# Disruption of the single-copy GOLDEN2-like gene underlies the classical *yellow* and *yellow-mutable* mutations of Japanese morning glory

**DOI:** 10.64898/2026.08.24.746626

**Authors:** Hibika Umehara, Kyoko Takagi, Soya Nakagawa, Shigeru Iida, Atsushi Hoshino

**Affiliations:** National Institute for Basic Biology, Okazaki 444-8585, Japan; Graduate Institute for Advanced Studies, The Graduate University for Advanced Studies, Okazaki 444-8585, Japan; Graduate School of Agriculture, Hokkaido University, Sapporo 060-8589, Japan; Institute of Science and Technology, Niigata University, Niigata 950-2181, Japan

**Keywords:** chloroplast development, Golden2-like (GLK), *Ipomoea nil*, transposable element, variegation

## Abstract

GOLDEN2-like (GLK) transcription factors are key regulators of chloroplast differentiation and photosynthetic gene expression. The classical *yellow* mutation in Japanese morning glory (*Ipomoea nil*) produces yellowish-green leaves, whereas an unstable allele, *yellow-mutable*, produces green somatic sectors on a yellowish-green background. The gene responsible for these mutations was identified as *InGLK*, which encodes a GOLDEN2-like transcription factor. The stable *yellow* mutant carried a 4-bp frameshift insertion in *InGLK*, whereas two *yellow-mutable* lines carried the *Tpn1*-family transposon *Tpn12* in intron 5. Excision of *Tpn12* in germinal revertants left short footprints and restored the green leaf phenotype. Genome searches identified InGLK as the sole GLK gene in *I. nil*.

Pigment analysis of green somatic reversion sectors and yellowish-green background areas showed that most of the measured photosynthetic pigments were significantly reduced in the yellowish-green background, whereas the chlorophyll a/b ratio was unchanged. Chloroplasts in the yellowish-green tissue retained thylakoid-like membranes and starch granule-like structures but had less distinct grana-like stacks and sparse stromal lamellae-like structures. Wild-type-like chloroplast ultrastructure was restored in germinal revertants. These findings show that loss of function of a single-copy GLK gene broadly reduces photosynthetic pigment accumulation and alters chloroplast internal membrane organization. The *yellow* mutants of *I. nil* therefore provide a genetic system for examining non-redundant GLK function.

## INTRODUCTION

The Japanese morning glory, *Ipomoea nil*, originated in tropical America and was introduced to Japan from China in the late eighth century as a medicinal plant (Hoshino et al. 2016). During the Edo period, extensive cultivation and selection by local enthusiasts led to the isolation of numerous spontaneous mutants, establishing a distinctive Japanese floricultural tradition centered on *Henka-Asagao*, ornamental morning glory mutants with diverse floral and leaf morphologies (Morita and Hoshino 2018). Many of these morphological changes affecting flowers and leaves are caused by the *Tpn1* family of *En/Spm*-like DNA transposons, which are the predominant endogenous mutagens in *I. nil* (Inagaki et al. 1994; Fukada-Tanaka et al. 2000; Morita et al. 2014; Morita et al. 2015; Hoshino et al. 2016). Together with the resources maintained by the National BioResource Project in Japan and the availability of a draft genome sequence, these mutant lines make *I. nil* a useful system for studying transposon-induced mutagenesis and gene regulation.

The *yellow* mutation is a classical leaf color mutation characterized by yellowish-green leaves (Fig. 1A) and has been used in genetic studies since the early twentieth century. In Japanese horticulture, this trait is known as “Kiba”, meaning yellow leaf, although the actual leaf color is yellowish-green rather than pure yellow. The *yellow* mutation has been widely retained in horticultural varieties. Although primarily recognized by its leaf phenotype, the mutation also affects other green aerial organs, including stems and sepals (Miyazawa 1929; Miyazawa 1932). Classical reports noted that yellow mutants show weaker vegetative growth than normal green plants (Imai 1927; Miyazawa 1929), and this reduced vigor has been considered useful in some horticultural contexts, where excessive vegetative growth can interfere with flower enlargement.

**Fig. 1.**
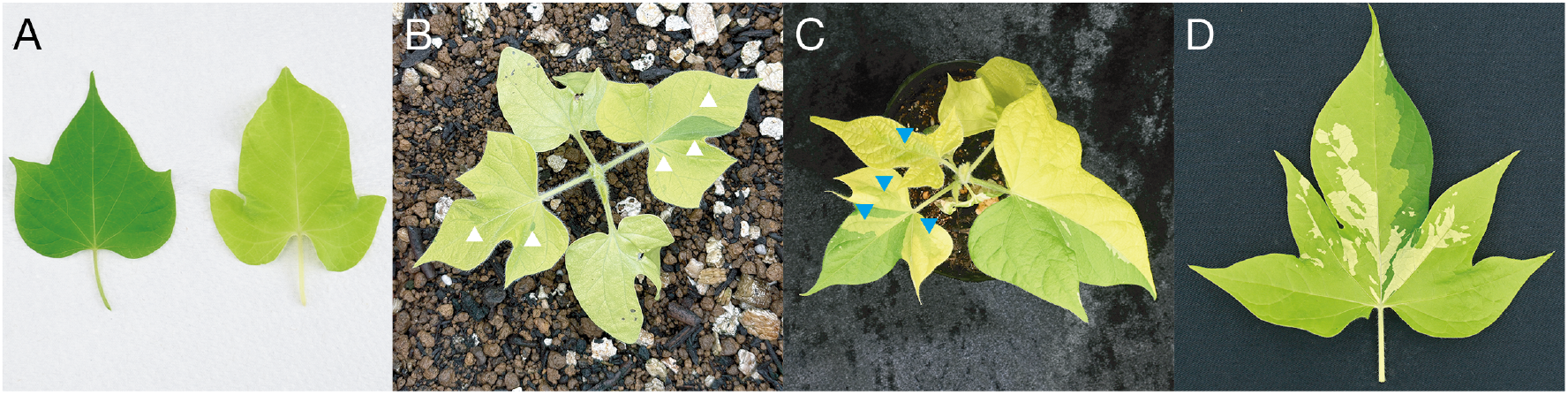
Leaf coloration and variegation phenotypes of stable *yellow* and *yellow-mutable* mutants. Representative plants or leaves are shown: (A) comparison of a wild-type leaf (TKS, left) and a stable *yellow* mutant leaf (AK36, right), (B) a *yellow-mutable* mutant (Q1075) exhibiting green somatic reversion sectors on a yellowish-green background, (C) a double mutant line (NIG1077) carrying both *yellow-mutable* and *variegated* mutations, and (D) a leaf from NIG1077. White arrowheads indicate green somatic reversion sectors in (B). Blue arrowheads indicate paler green sectors in (C), which may reflect somatic reversion events affecting internal leaf tissue. In (C) and (D), white to pale-yellow sectors are caused by the *variegated* mutation.

An unstable allele of *yellow*, referred to as *yellow-mutable*, produces green somatic sectors on a yellowish-green background (Fig. 1B–D). This pattern is genetically distinct from the *variegated* mutation, which is controlled by a separate locus and produces white or yellow variegation on a green background (Imai 1927; Miyazawa 1929; Imai 1930a). The *yellow-mutable* phenotype is historically called “Matsushima”, after Matsushima, one of the Three Views of Japan, where pine-covered islands are scattered across the sea. The green spots on yellowish-green leaves were likened to these islands. Imai (1927) concluded from a survey of early horticultural records that many old “yellow” leaves were not ordinary stable *yellow* forms, but Matsushima-lined yellow forms that produced green sectors or green-variegated yellow leaves. He further noted that Matsushima-lined yellows became rare after the Meiji era and were gradually replaced by ordinary *yellow* forms that bred true. Subsequent classical studies established an allelic relationship in which the wild-type green allele is dominant to *yellow-mutable*, which in turn is dominant to stable *yellow* with respect to the production of green sectors (Miyazawa 1929; Imai 1930b; Miyazawa 1932). These studies also proposed that green sectors arise through somatic reversion of the *yellow* factor to the green state. This interpretation is consistent with the behavior of transposon-induced mutable alleles, but the responsible gene and molecular basis of the *yellow-mutable* phenotype remained unknown.

GOLDEN2-like (GLK) transcription factors are key regulators of chloroplast differentiation and nuclear photosynthetic gene expression in land plants. GLK proteins activate genes involved in light-harvesting complex formation, chlorophyll biosynthesis, and other chloroplast-related processes (Hall et al. 1998; Fitter et al. 2002; Waters et al. 2009; Chen et al. 2016; Choi et al. 2024; Yelina et al. 2024). In addition to their central roles in chloroplast development, GLK pathways have also been implicated in diverse processes, including stress responses, stomatal regulation, disease resistance, and specialized metabolism (Savitch et al. 2007; Nagatoshi et al. 2016; Ahmad et al. 2019; Wang et al. 2022). However, the phenotypic consequences of GLK loss differ among species and organs, largely because of variation in gene copy number, functional redundancy, and tissue specificity.

The classical maize *golden2* mutant first demonstrated the requirement of GLK activity for chloroplast development, particularly in bundle sheath cells of C4 leaves, and maize possesses two GLK genes with partially specialized functions in bundle sheath and mesophyll cells (Hall et al. 1998; Cribb et al. 2001; Rossini et al. 2001). In many C3 model plants, GLK genes occur as paralogous pairs. In Arabidopsis and rice, single *glk* mutants show weak or limited leaf phenotypes because of functional compensation, whereas double mutants exhibit severe chlorosis (Fitter et al. 2002; Waters et al. 2009). Similar redundancy has been reported in the moss *Physcomitrium patens*, in which double mutants of *PpGLK1* and *PpGLK2* show pale-green phenotypes (Yasumura et al. 2005). In contrast, the liverwort *Marchantia polymorpha* possesses a single *GLK* gene, and loss-of-function mutants have revealed an essential role of GLK in chloroplast biogenesis in an early-diverging land plant lineage (Yelina et al. 2024). Loss-of-function studies in crops further indicate that GLK function can be strongly tissue specific. In tomato, *SlGLK2* corresponds to the *uniform ripening* locus and regulates chloroplast development in fruit (Powell et al. 2012; Nguyen et al. 2014), whereas in barley, *HvGLK2* mutations cause chlorophyll-deficient hulls while leaf blades remain largely green, presumably because of compensation by *HvGLK1* (Taketa et al. 2021). Thus, although GLK function in plastid development is broadly conserved, its redundancy and tissue specificity vary substantially among plant lineages.

In this study, the gene responsible for the *yellow* mutation was identified by transposon display as an *I. nil* homolog of GLK, designated *InGLK*. The structures of the stable *yellow* and *yellow-mutable* alleles and the excision footprints in germinal revertants were characterized, and the effects of *InGLK* deficiency on photosynthetic pigment accumulation and chloroplast ultrastructure were examined. Green somatic reversion sectors and adjacent yellowish-green background areas of the *yellow-mutable* line were also used to explore the association between local *InGLK* status and florigen gene expression.

## RESULTS

### Identification of the *InGLK* gene and allelic structures

Representative phenotypes of the stable *yellow* and *yellow-mutable* mutants are shown in Fig. 1. The transposon display method utilizing the *Tpn1* family of transposons, the primary mutagens in *I. nil*, has successfully identified several genes responsible for mutant phenotypes (Fukada-Tanaka et al. 2000; Fukada-Tanaka et al. 2001; Iwasaki and Nitasaka 2006; Morita et al. 2014). Therefore, the gene responsible for the *yellow* mutation was isolated using this approach. First, germinal revertants with wild-type green leaves were obtained from two *yellow-mutable* lines, NIG553 and K41. Analysis of phenotypic segregation in the subsequent generation revealed that two individuals derived from line NIG553 and three from line K41 were homozygous for revertant alleles. Consequently, genomic DNA was extracted and compared between these five germinal revertants and eight individuals showing the *yellow-mutable* phenotype from each line as controls (Supplementary Fig. S1A). This analysis identified a 116-bp sequence that completely co-segregated with the mutant phenotype. Sequence analysis revealed that this fragment contained a portion of intron 5 and exon 6 of the *InGLK* gene (LOC109188258). Subsequently, full-length sequencing of *InGLK* amplified by PCR from the two mutant lines demonstrated that a *Tpn1*-family transposon, designated *Tpn12*, was inserted adjacent to the sequence identified by transposon display, specifically 14 bp upstream of exon 6 (Fig. 2A, B, Supplementary Fig. S1B). Furthermore, all analyzed germinal revertants retained a 3- to 5-bp footprint sequence as a scar of transposon excision (Fig. 2B). A 5-bp footprint sequence was also identified in a homozygous germinal revertant isolated from another *yellow-mutable* line, NIG1073. These results demonstrate that the gene responsible for the *yellow* mutation is *InGLK*.

**Fig. 2.**
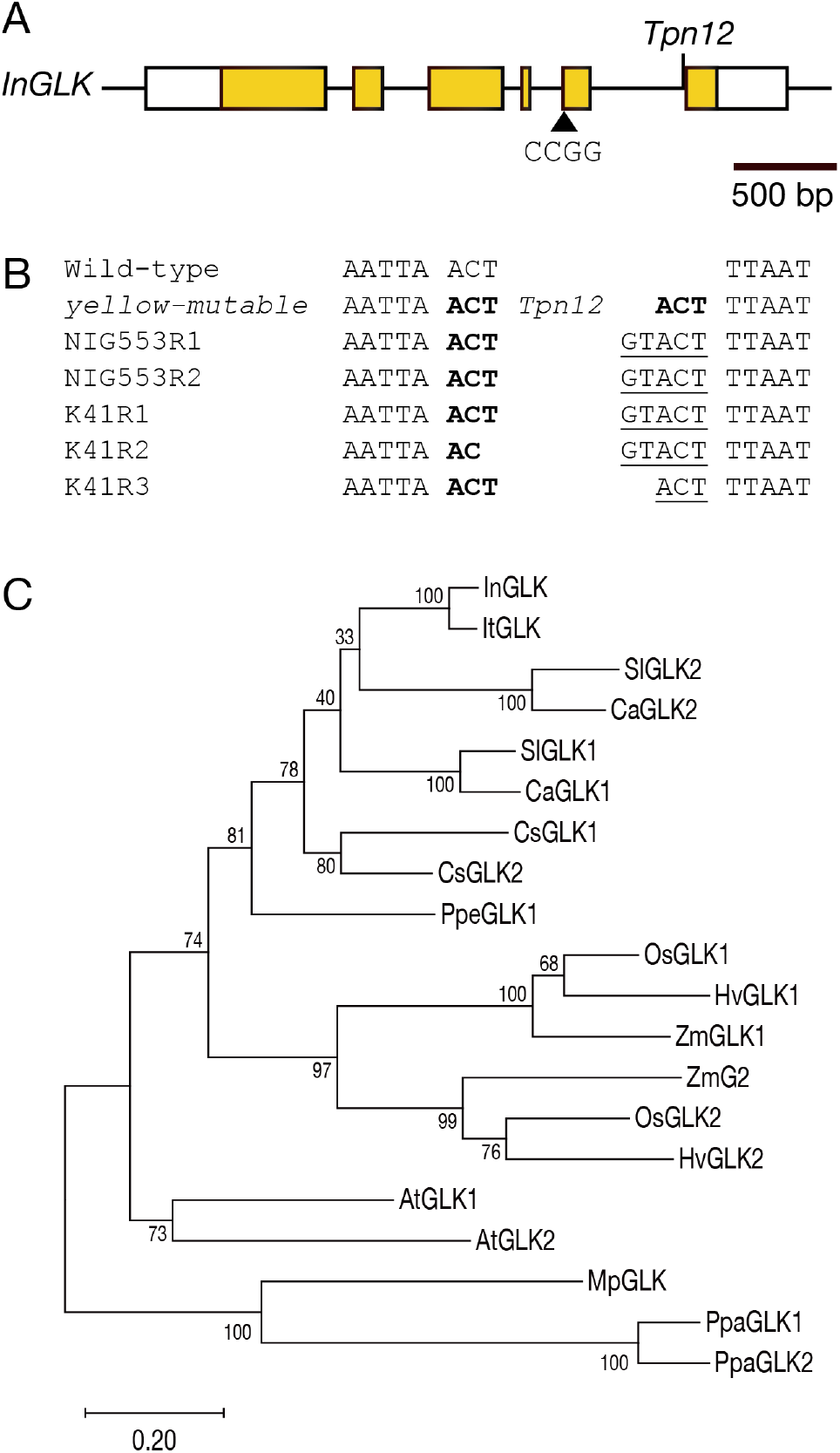
The single-copy *InGLK* gene is disrupted by a transposon or a short insertion in the *yellow* mutants. (A) Schematic representation of the *InGLK* gene structure and the precise mutation sites. Exons are depicted as boxes and introns as lines. Yellow and white boxes indicate coding and untranslated regions, respectively. The *yellow-mutable* allele harbors a *Tpn12* transposon insertion within intron 5, whereas the stable *yellow* allele contains a 4-bp insertion in exon 5. (B) Sequence alignments showing the footprint sequences generated by germinal excision of *Tpn12*. The 3-bp target site duplication is shown in bold, and footprint sequences are underlined. (C) Phylogenetic analysis of GLK proteins showing that InGLK belongs to the GLK clade. The tree was constructed using the amino acid sequences listed in Supplementary Table S3. Bootstrap values were calculated from 1,000 replicates and are shown at the nodes. Species abbreviations are as follows: In, *Ipomoea nil*; It, *Ipomoea triloba*; Sl, *Solanum lycopersicum*; Ca, *Capsicum annuum*; Cs, *Camellia sinensis*; Ppe, *Prunus persica*; Os, *Oryza sativa*; Hv, *Hordeum vulgare*; Zm, *Zea mays*; At, *Arabidopsis thaliana*; Mp, *Marchantia polymorpha*; Ppa, *Physcomitrium patens*.

In the initial genome annotation of *I. nil* (Hoshino et al. 2016), this locus was erroneously annotated as two separate genes: INIL03g17681, corresponding to the 5′ region, and INIL03g17682, corresponding to the 3′ region. The gene is located on chromosome 3 at positions 33,587,964–33,591,083. Classical genetic analysis had previously established that the *yellow* locus is tightly linked to the *dusky* locus, with a map distance of 1.2 cM (Imai and Tabuchi 1934; Hagiwara 1956). Consistent with these classical genetic data, the physical map revealed that *InGLK* is located only 220 kb away from the *Dusky* gene, LOC109188143/INIL03g17665, which is located on chromosome 3 at positions 33,365,935–33,367,605. *Dusky* was previously identified as *InUGT78D2*, encoding UDP-glucose:anthocyanidin 3-*O*-glucoside-2′′-*O*-glucosyltransferase, by Morita et al. (2005).

RT-PCR failed to amplify exon 6-containing *InGLK* cDNA from the *yellow-mutable* mutant lines, whereas amplification was detected in wild-type plants and germinal revertant plants, suggesting that *yellow-mutable* is a severe loss-of-function allele (Supplementary Fig. S1C).

Meanwhile, full-length sequence analysis of *InGLK* in the stable *yellow* mutant line AK55 revealed a 4-bp insertion of “CCGG” in exon 5, which causes a frameshift mutation (Fig. 2A). Furthermore, examination of an additional 13 *yellow* mutant lines showed that all lines carried the same 4-bp insertion, whereas this insertion was absent from all 21 wild-type lines examined (Supplementary Table S1). Notably, this 4-bp insertion sequence did not contain the 3-bp target site duplication (TSD) characteristic of *Tpn1*-family transposon footprints.

A BLASTP search against the Japanese morning glory genome database identified only one GLK homolog, InGLK. An NCBI BLASTP search restricted to the genus *Ipomoea* identified a single annotated GLK homolog in *I. triloba*, designated ItGLK. Phylogenetic analysis placed InGLK and ItGLK within the GLK clade (Fig. 2C).

### Expression pattern of *InGLK* in different tissues

To examine the tissue expression pattern of *InGLK*, publicly available RNA-seq data from six tissues were analyzed. Based on TPM values, *InGLK* showed the highest expression in leaves, followed by stems and embryos (Table 1). Lower expression was observed in flowers and roots, and only marginal expression was detected in seed coats (Table 1). Because these data were derived from single RNA-seq samples for each tissue, statistical testing was not performed.

**Table 1.**
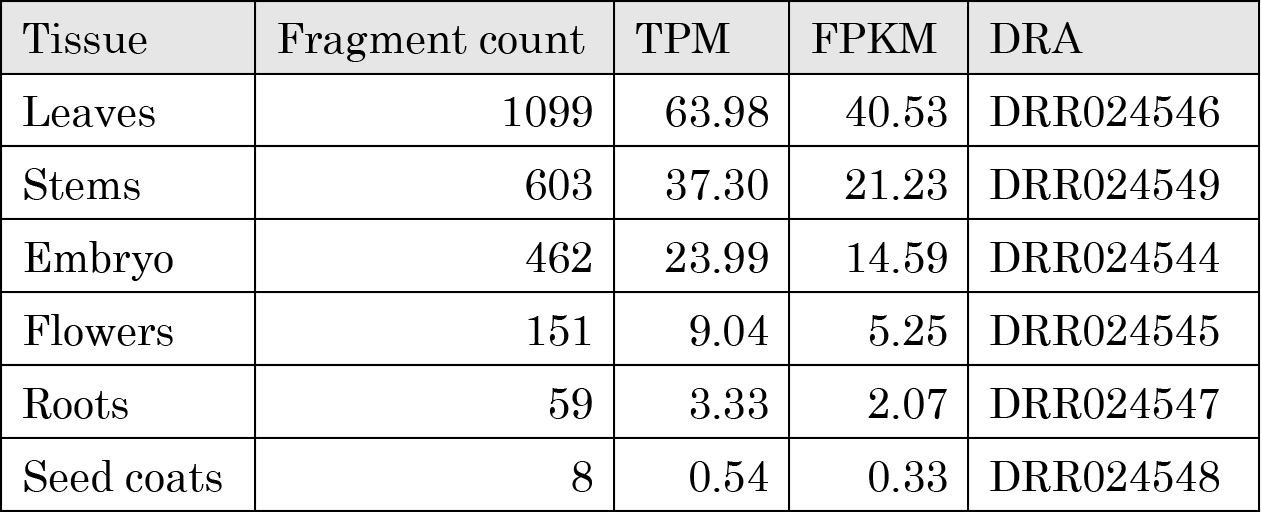
Expression levels of *InGLK* in six tissues based on public RNA-seq data. Fragment count indicates the number of paired-end fragments assigned to *InGLK*. TPM, transcripts per million; FPKM, fragments per kilobase of transcript per million mapped reads. No statistical tests were performed because biological replicates were unavailable.

### Photosynthetic pigment accumulation

To verify whether the leaf color change in the yellowish-green background of the *yellow-mutable* lines reflects a decrease in pigment content, quantitative analysis was performed using Ultra-Performance Liquid Chromatography (UPLC). Yellowish-green background areas and green somatic reversion sectors were excised from true leaves of the Q1075 line. Eight major photosynthetic pigments were targeted for measurement: chlorophyll a, chlorophyll b, β-carotene, lutein, neoxanthin, violaxanthin, antheraxanthin, and zeaxanthin.

The analysis revealed that the contents of most measured pigments were significantly reduced in the yellowish-green background compared with the green sectors (Fig. 3). Although antheraxanthin showed a decreasing trend, the difference was not statistically significant.

**Fig. 3.**
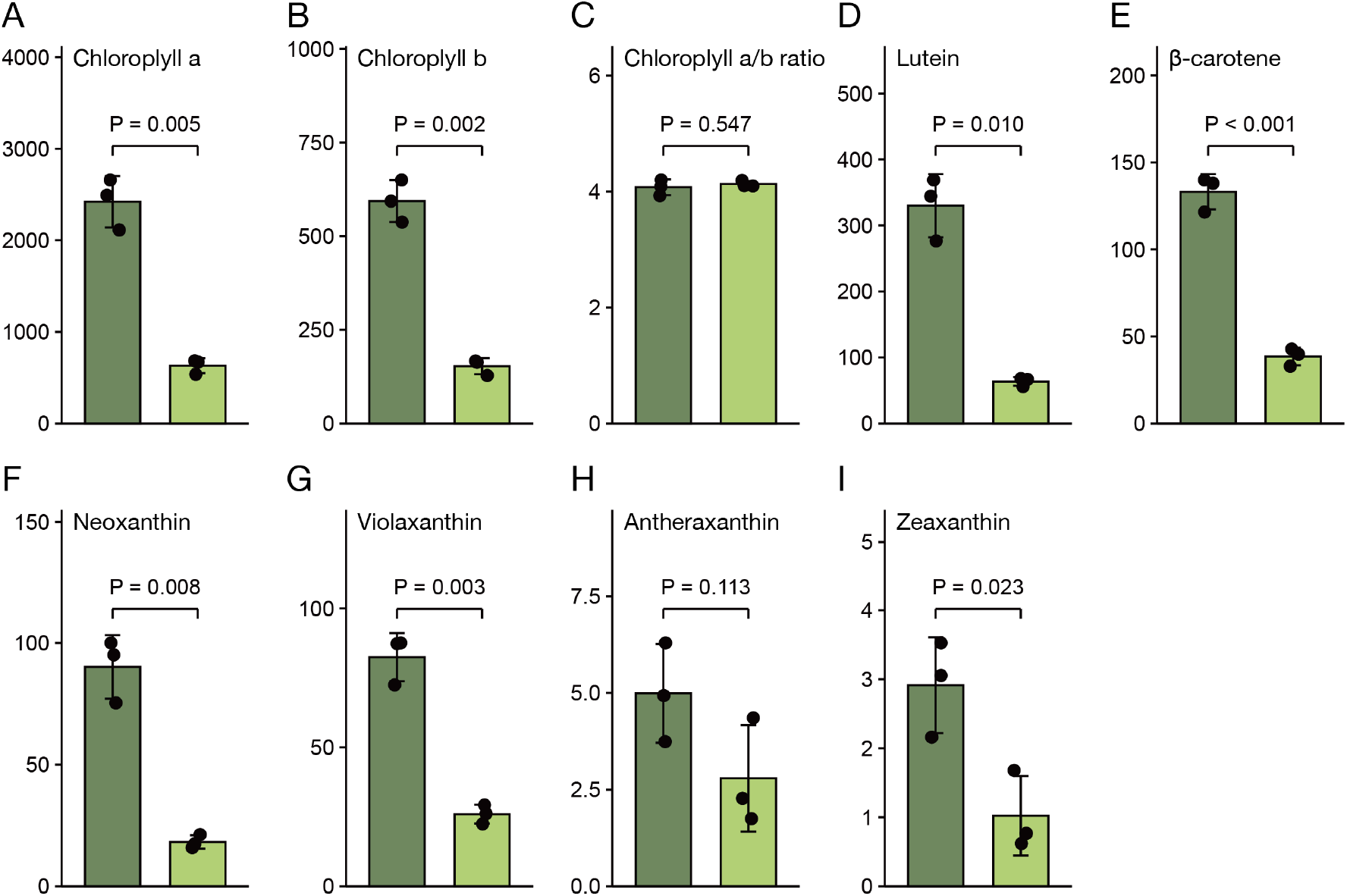
Pigment contents in green somatic reversion sectors and yellowish-green background areas of yellow-mutable leaves. The contents of chlorophyll a (A), chlorophyll b (B), lutein (D), β-carotene (E), neoxanthin (F), violaxanthin (G), antheraxanthin (H), and zeaxanthin (I) were quantified by UPLC, and the chlorophyll a/b ratio was calculated (C). Values are shown as pigment contents (µg g⁻¹ FW), except for the chlorophyll a/b ratio. Dark green left bars and light green right bars indicate green reversion sectors and yellowish-green background areas, respectively. Data represent mean ± SD of three biological replicates. *P* values were calculated using Welch’s *t*-test.

Furthermore, the chlorophyll a/b ratio was calculated to examine whether the relative composition of the chlorophyll system differed between the two sectors. No significant difference was detected between the green reversion sectors and the yellowish-green background areas (p = 0.547).

These results indicate that the yellowish-green phenotype is associated with a broad reduction in photosynthetic pigment accumulation rather than a selective reduction in a particular pigment class.

### Chloroplast ultrastructure

The chloroplast ultrastructure in leaves of the wild-type, the *yellow-mutable* line, and its germinal revertant was examined using transmission electron microscopy (Fig. 4). In the wild-type, well-developed chloroplasts were observed in mesophyll cells (Fig. 4A, B). Within these chloroplasts, thylakoid membranes, grana-like stacked structures, and stromal lamellae-like structures were clearly recognized. Large structures with low electron density were also observed within the chloroplasts and were considered to be starch granules.

**Fig. 4.**
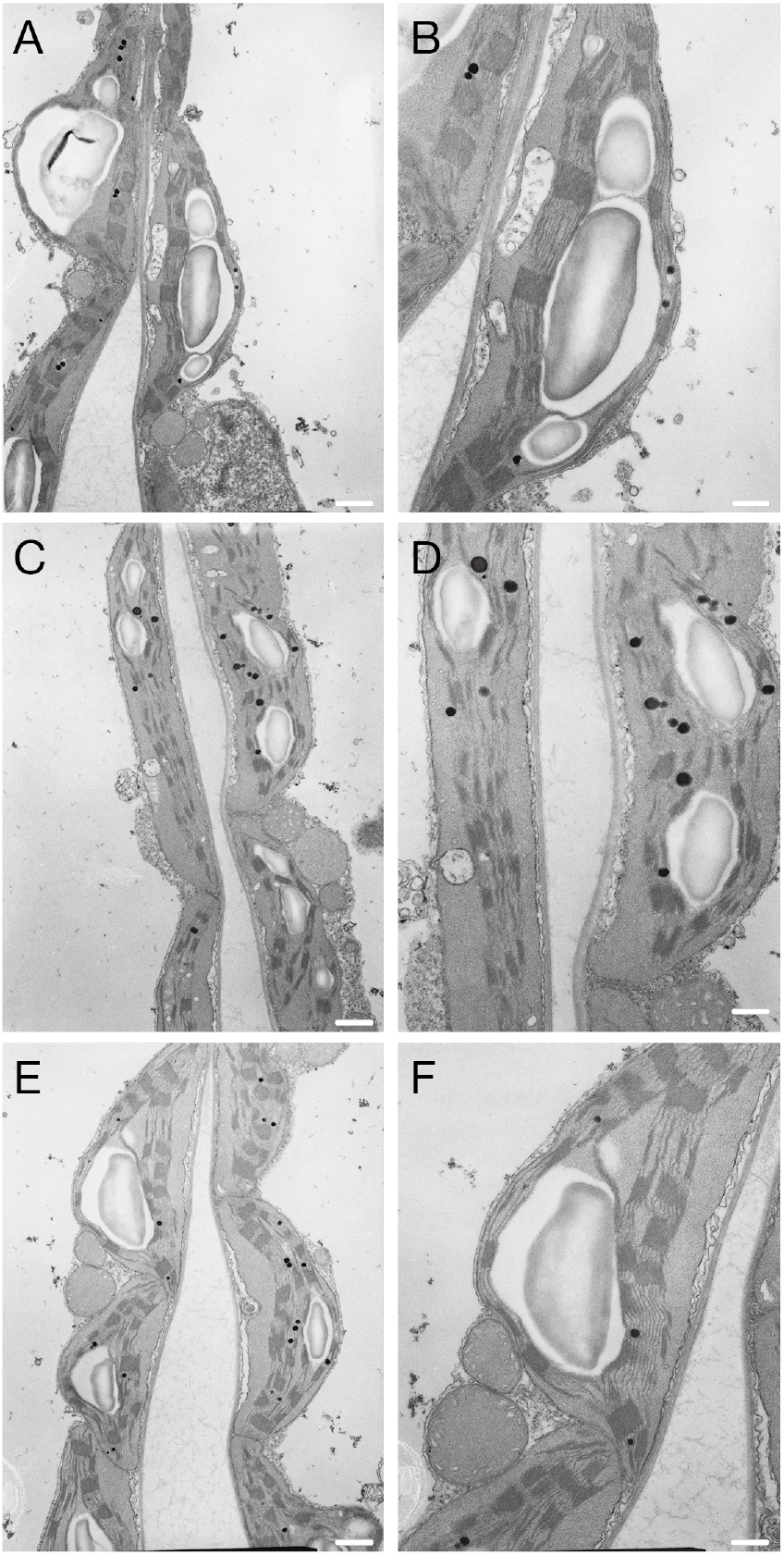
Chloroplast ultrastructure in mesophyll cells of wild-type, *yellow-mutable*, and germinal revertant leaves. Transmission electron microscopy was used to compare chloroplasts in mesophyll cells of wild-type plants (A, B), the yellowish-green background tissue of *yellow-mutable* leaves (C, D), and germinal revertant plants (E, F). Panels B, D, and F show enlarged views of A, C, and E, respectively. Scale bars: 1 µm in A, C, and E; 500 nm in B, D, and F.

Chloroplasts were also observed in mesophyll cells from the yellowish-green background tissue of the *yellow-mutable* line (Fig. 4C, D), and internal membrane structures resembling thylakoid membranes were recognized. However, compared with the wild-type, the grana-like stacked structures were less distinct, and the stromal lamellae-like structures were sparse. Nevertheless, structures with low electron density, resembling starch granules, were also observed in the chloroplasts of the mutant.

In the germinal revertant, thylakoid membranes, grana-like stacked structures, and stromal lamellae-like structures were clearly recognized within the chloroplasts (Fig. 4E, F). Structures with low electron density, resembling starch granules, were also observed. The chloroplast ultrastructure of the revertant was morphologically closer to that of the wild-type than to that of the mutant, indicating that the alterations in the chloroplast internal membrane structure observed in the mutant were restored to a wild-type-like state in the revertant.

An internally controlled comparison was also performed using a single transverse section of a *yellow-mutable* leaf containing both green somatic reversion and yellowish-green background tissues. In the available images of the spongy mesophyll, grana-like stacks in the green somatic reversion sector appeared thicker and more distinct than those in the yellowish-green background area. In contrast, the limited images obtained from the palisade mesophyll did not allow a clear sector-associated comparison (Supplementary Fig. S2).

These results indicate that *InGLK* deficiency alters chloroplast internal membrane organization, particularly the appearance of grana-like stacks and stromal lamellae-like structures, while leaving starch granule-like structures detectable.

### Association between *InGLK* status and *InFT1* and *InFT2* expression

To explore whether local *InGLK* status is associated with the expression of FT homologs, RT-qPCR analysis was performed for *InFT1* (INIL09g31483) and *InFT2* (INIL03g17943). These genes were previously characterized as *PnFT1* and *PnFT2*, respectively, in *Pharbitis nil*, the former name of *I. nil* (Hayama et al. 2007). The *yellow-mutable* line produces green somatic reversion sectors on a yellowish-green background, allowing paired comparisons of tissues with different *InGLK* status within the same leaf. Based on the reported late-night increase in *PnFT1* and *PnFT2* expression under inductive short-day conditions (Hayama et al. 2007), sampling was performed approximately 5 min after light-on, following an 11-h dark period. Green reversion sectors and corresponding yellowish-green background areas were collected from three true leaves of different ages and positions from a single individual. Each leaf was divided along the midrib to obtain internally controlled paired samples (Supplementary Fig. S3A).

In all three leaves, transcript levels of both *InFT1* and *InFT2* were lower in the yellowish-green background areas than in the corresponding green reversion sectors (Supplementary Fig. S3B). The difference was more pronounced for *InFT2*, whereas *InFT1* showed a smaller but consistent reduction. Because the three leaves were obtained from a single individual and differed in age and position, they were analyzed as individual paired samples rather than as independent biological replicates. Nevertheless, the consistent within-leaf pattern suggests an association between local *InGLK* status and the expression of *InFT1* and *InFT2*.

## DISCUSSION

### A single-copy *GLK* gene underlies the classical *yellow* mutation in *I. nil*

This study identified *Yellow* as encoding a GOLDEN2-like transcription factor, designated *InGLK*. The structures of the stable *yellow* and *yellow-mutable* alleles and the excision footprints found in germinal revertants support this conclusion. The stable *yellow* mutant carries a frameshift insertion in *InGLK*, whereas the *yellow-mutable* allele contains the novel *Tpn1*-family transposon *Tpn12* in intron 5. In germinal revertants derived from *yellow-mutable* mutants, excision of *Tpn12* left short footprints and was associated with restoration of the green leaf phenotype. These results provide a molecular explanation for classical genetic observations that green sectors in *yellow-mutable* leaves arise through somatic or germinal reversion of the *yellow* factor. The contrasting allele structures are also consistent with the historical distinction between Matsushima-lined yellows and ordinary yellow forms that bred true (Imai 1927).

Genome searches identified InGLK as the sole GLK gene in the *I. nil* genome. This feature distinguishes *I. nil* from many model angiosperms in which duplicated *GLK* genes partially compensate for each other (Wang et al. 2013; Chen et al. 2016). In Arabidopsis and rice, single *glk* mutants generally show weak or limited leaf phenotypes, whereas double mutants exhibit stronger chlorotic phenotypes (Fitter et al. 2002; Waters et al. 2009; Wang et al. 2013). In contrast, disruption of the single *InGLK* gene in *I. nil* produces a systemic yellowish-green leaf phenotype. Thus, *I. nil* provides a useful system for examining non-redundant GLK function in leaves.

The systemic phenotype of stable *yellow* mutants is consistent with classical descriptions of their weaker vegetative growth compared with normal green plants (Imai 1927; Miyazawa 1929). Photosynthetic activity was not directly measured in the present study. Nevertheless, the marked reduction in photosynthetic pigment accumulation and the alteration of chloroplast internal membrane structure provide a cellular basis for the yellowish-green phenotype and may be related to the reduced vigor described in earlier studies.

### *InGLK* deficiency reduces pigment accumulation and alters chloroplast ultrastructure

Quantitative pigment analysis showed that most major photosynthetic pigments, including chlorophylls and carotenoids, were significantly reduced in the yellowish-green background of *yellow-mutable* leaves compared with green reversion sectors. This result indicates that *InGLK* is required for normal accumulation of photosynthetic pigments in *I. nil* leaves. However, the chlorophyll a/b ratio did not differ significantly between the yellowish-green background and green sectors. These findings suggest that InGLK deficiency broadly reduces photosynthetic pigment accumulation without causing a detectable shift in the relative proportions of chlorophyll a and chlorophyll b.

TEM observations also revealed alterations in chloroplast ultrastructure. Chloroplasts in the yellowish-green background retained recognizable thylakoid-like membranes and starch granule-like structures, indicating that chloroplast development was not completely blocked. Nevertheless, compared with wild-type and germinal revertant chloroplasts, grana-like stacked structures were less distinct and stromal lamellae-like structures were sparse. These observations indicate that *InGLK* is required for normal organization of the chloroplast internal membrane system. The recovery of wild-type-like chloroplast ultrastructure in germinal revertants further supports the association between these ultrastructural alterations and disruption of *InGLK*.

The phenotype of the *I. nil yellow* mutant can be compared with *glk* mutants in other plants. In Arabidopsis and rice, strong chlorotic phenotypes are most evident when redundant GLK functions are disrupted (Fitter et al. 2002; Waters et al. 2009; Wang et al. 2013). The classical maize *golden2* mutant affects chloroplast differentiation in a cell-type-specific manner, particularly in bundle sheath cells (Hall et al. 1998; Cribb et al. 2001; Rossini et al. 2001; Lambret-Frotte et al. 2024). In the liverwort *Marchantia polymorpha*, which possesses a single *GLK* gene, loss-of-function mutants show defective chloroplast biogenesis (Yelina et al. 2024). Crop studies also indicate that GLK function can be tissue specific: tomato *SlGLK2* controls fruit chloroplast development and underlies the *uniform ripening* phenotype (Powell et al. 2012; Nguyen et al. 2014), whereas barley *HvGLK2* mutations affect chlorophyll accumulation in hulls while leaves remain largely green, presumably because of compensation by *HvGLK1* (Taketa et al. 2021).

Compared with these examples, yellowish-green tissues of *I. nil* retained recognizable thylakoid-like membranes and did not show a significant change in the chlorophyll *a/b* ratio. This suggests that basal chloroplast development can proceed to some extent even when *InGLK* function is impaired. Thus, *InGLK* appears to act primarily as a quantitative regulator of chloroplast development in leaves, promoting normal levels of pigment accumulation and internal membrane organization rather than serving as an all-or-none determinant of chloroplast formation. Further analysis of photosynthetic protein accumulation, chloroplast gene expression, and direct photosynthetic activity will be needed to define the molecular and physiological effects of *InGLK* deficiency more precisely.

### Association between InGLK-dependent chloroplast development and florigen gene expression

The *yellow-mutable* system provides a distinctive opportunity to examine local effects associated with *InGLK* function, because green somatic reversion sectors and yellowish-green mutant tissue occur within the same leaf. This paired-sector design reduces differences in genetic background, growth conditions, sampling time, and systemic developmental status. In all three leaves examined, transcript levels of both *InFT1* and *InFT2* were lower in the yellowish-green background areas than in the corresponding green reversion sectors (Supplementary Fig. S3). Although the leaves differed in age and position, they were obtained from a single plant and therefore do not represent independent biological replicates. Nevertheless, the consistent within-leaf pattern suggests a local association between *InGLK*-dependent chloroplast development and florigen gene expression, rather than demonstrating direct regulation of *InFT* expression by *InGLK*.

Reduced *InFT1* and *InFT2* expression does not necessarily indicate a major effect of *InGLK* on flowering time. Stable *yellow* mutants have not been reported to show an obvious flowering-time phenotype in classical descriptions, and no conspicuous difference has been noticed during routine cultivation of multiple *yellow* mutant lines. However, flowering time was not quantitatively analyzed in the present study. In *Arabidopsis*, *GLK1* and *GLK2* promote *BBX14*, *BBX15*, and *BBX16* expression, thereby repressing *CO*-mediated *FT* expression and precocious flowering (Susila et al. 2023), whereas expression of *InFT1* and *InFT2* was lower in *InGLK*-deficient tissue of *I. nil*. The tissue-specific phenotypes of barley *HvGLK2* mutants (Taketa et al. 2021) and the recently demonstrated joint regulation of chloroplast development by GLK and B-box proteins in *Arabidopsis* (Kakuda et al. 2026) further indicate that GLK-associated phenotypes and downstream responses are context dependent. Thus, the relationship among GLK activity, chloroplast development, *FT* expression, and flowering may depend on species, tissue, and photoperiodic context.

Overall, the structures of the stable *yellow* and *yellow-mutable* alleles, together with the excision footprints found in germinal revertants, establish *InGLK*, the sole GLK gene in *I. nil*, as the gene responsible for the classical *yellow* mutations. The *yellow* mutants of *I. nil* provide a useful system for examining the non-redundant role of a single-copy GLK gene in photosynthetic pigment accumulation and chloroplast internal membrane organization.

## MATERIALS AND METHODS

### Plant material

The plants used in this study are listed in Supplementary Table S1. Plants used for electron microscopy, pigment analysis, and RT-qPCR were grown in an air-conditioned growth room on racks equipped with fluorescent lamps under a 13-h light/11-h dark cycle. The other plants were cultivated in a greenhouse. Genomic DNA was extracted using either the Genomic-tip 500/G (QIAGEN, Hilden, Germany) or the GENE PREP STAR PI-480 (KURABO, Osaka, Japan), as previously described (Hoshino et al. 2016).

### Isolation of the *Yellow* gene

The simplified transposon display method (Fukada-Tanaka et al. 2001; Morita et al. 2014) was used with minor modifications to clone the gene responsible for the *yellow-mutable* phenotype. Genomic DNA was digested with the restriction enzyme *Bfa*I and ligated to specific adapters (Supplementary Table S2). PCR was performed using adapter-specific primers and primers specific to the terminal sequences of *Tpn1*-family transposons. Bands co-segregating with the *yellow-mutable* phenotype were isolated and sequenced. To examine the allelic structures of the *Yellow* locus, PCR was performed using primers Matsushima-Fw4 and Matsushima-Rv6. Because the *yellow-mutable* alleles contained the *Tpn12* insertion and were not efficiently amplified with this primer pair, the insertion junctions were analyzed using primer combinations Matsushima-Fw5/TIR-OUT-S and Matsushima-Rv6/TIR-OUT-S.

### Full-length sequence analysis using nanopore sequencing

To characterize the *yellow* alleles, long-range PCR was performed using PrimeSTAR GXL DNA Polymerase (Takara Bio, Kusatsu, Japan) with primers Matsushima-Fw10 and Matsushima-Rv17. Genomic DNA samples from the *yellow-mutable* lines NIG553 and K41 and the stable *yellow* line AK55 were used as templates. The PCR products were sequenced using the Plasmid-EZ service (Azenta Life Sciences, Tokyo, Japan), which utilizes nanopore sequencing technology, following the method described by Nakagawa et al. (2025b). For line K41, however, the PCR product was subjected to nanopore sequencing without the circularization step described in the original method.

### RT-PCR analysis

Total RNA was isolated from leaves by guanidinium thiocyanate–cesium chloride ultracentrifugation as described previously (Abe et al. 1997). First-strand cDNA was synthesized using SuperScript III Reverse Transcriptase (Thermo Fisher Scientific, Waltham, MA, USA), and RT-PCR was performed using TaKaRa Ex Taq DNA Polymerase (Takara Bio, Kusatsu, Japan). *GAPDH2* (INIL15g35608) transcripts were amplified as an internal control. Primer sequences are listed in Supplementary Table S2.

### Identification of GLK homologs

GLK homologs were searched by BLASTP against the Japanese morning glory genome database and the NCBI non-redundant protein database restricted to the genus *Ipomoea*, using Arabidopsis GLK1 (AtGLK1; AT2G20570) as the query. To assess whether an additional GLK homolog was present in *I. nil*, the second-highest-scoring *I. nil* protein was subjected to a reciprocal BLASTP search against Arabidopsis proteins.

### Phylogenetic analysis

GLK amino acid sequences listed in Supplementary Table S3 were aligned using MUSCLE with default parameters in MEGA12 (Edgar 2004; Kumar et al. 2024). A maximum likelihood phylogenetic tree was constructed in MEGA12 using the JTT amino acid substitution model with uniform rates among sites (Jones et al. 1992; Kumar et al. 2024). Gaps and missing data were treated by partial deletion with a site coverage cutoff of 80%. The nearest-neighbor-interchange method was used for the ML heuristic search. Branch support was evaluated by bootstrap analysis with 1,000 replicates.

### RNA-seq reanalysis of public transcriptome data

Publicly available paired-end RNA-seq reads from six tissues of *I. nil* were reanalyzed to examine the tissue expression pattern of *InGLK*. The six datasets, DRR024544–DRR024549, were mapped to the *I. nil* Asagao_1.1 reference genome sequence, GCF_001879475.1_Asagao_1.1_genomic.fna, using HISAT2 v2.2.1. The resulting SAM files were converted to sorted BAM files and indexed using SAMtools v1.21. Gene-level TPM and FPKM values were estimated using StringTie v3.0.0 with the -e option and the corresponding RefSeq GFF annotation file from which organellar sequences had been removed. Gene-level fragment counts were calculated separately using featureCounts in the Subread package v1.6.1 with the options -p, -t exon, and -g gene. The resulting gene-level tables of fragment counts, TPM values, and FPKM values are provided as Supplementary Tables S4, S5, and S6, respectively, and were used to extract the expression values for *InGLK*.

### Pigment analysis

Leaf discs were collected from the green somatic reversion sectors and yellowish-green background areas of *yellow-mutable* leaves. Frozen *I. nil* leaf discs (0.15 g) were homogenized in liquid nitrogen using a mortar and pestle, and pigments were extracted with 1 mL of acetone. After centrifugation at 15,000 rpm for 5 min, the supernatants were subjected to pigment analysis. Pigment analysis was performed using an ACQUITY UPLC system (Waters, Milford, MA, USA) with the same chromatographic settings as described by Aihara et al. (2019). The following eight pigments were separated and quantified: neoxanthin, violaxanthin, antheraxanthin, zeaxanthin, lutein, chlorophyll *b*, chlorophyll *a*, and β-carotene. Pigment standards were purchased from DHI Laboratory Products (Hørsholm, Denmark). Chlorophyll a/b ratios were calculated separately for each biological replicate. Differences between green reversion sectors and yellowish-green background areas were evaluated using Welch’s *t*-test in R version 4.3.2.

### Ultrastructure analysis

To examine chloroplast ultrastructure, leaf tissues were collected from the TKS wild type, the yellowish-green K41 mutant, and its germinal revertant. For an internally controlled comparison, a transverse section spanning the midrib was prepared from a Q1072 leaf, in which the two sides of the midrib comprised yellowish-green background tissue and green somatic reversion tissue, respectively. Sample fixation, resin embedding, and transmission electron microscopy were performed by Hanaichi Ultrastructure Research Institute (Okazaki, Japan). Samples from TKS, K41, and the K41 germinal revertant were observed using a JEM-1200EX transmission electron microscope, whereas the Q1072 sample was observed using a JEM-1400Flash transmission electron microscope (JEOL, Tokyo, Japan).

### RT-qPCR for *InFT* gene expression

Line Q1075 was grown under long-day conditions in the growth room described above, with lights on at 6:15 and off at 19:15. Three true leaves of different ages and positions were sampled simultaneously from a single individual (plant ID: 2024319). Each leaf was divided along the midrib into a green reversion sector and the corresponding yellowish-green background area at approximately 6:20, about 5 min after light-on, to obtain internally controlled paired samples.

Total RNA was extracted using the Maxwell RSC Plant RNA Kit (Promega, Madison, WI, USA), and cDNA was synthesized using ReverTra Ace qPCR RT Master Mix with gDNA Remover (TOYOBO, Osaka, Japan). The expression of the *FT* homologs *InFT1* and *InFT2* was analyzed by quantitative PCR using THUNDERBIRD Next SYBR qPCR Mix (TOYOBO, Osaka, Japan) and a QuantStudio 3 Real-Time PCR System (Thermo Fisher Scientific, Waltham, MA, USA). Primer sequences are listed in Supplementary Table S2. Expression levels were calculated from Quantity Mean values obtained using standard curves generated from serial dilutions of plasmid DNA or cDNA-derived fragments and normalized to *gATPase* (INIL09g35921; Nakagawa et al. 2025a). Technical triplicates were averaged for each sample. The three paired leaf samples were analyzed individually to retain variation associated with leaf age or position.

## DECLARATIONS

### Data availability

The nucleotide sequences of the *InGLK* alleles determined in this study have been deposited in the DNA Data Bank of Japan (DDBJ) under accession numbers LC942611 for the NIG553 *yellow-mutable* allele, LC942612 for the K41 *yellow-mutable* allele, and LC942613 for the AK55 stable *yellow* allele. The public RNA-seq datasets reanalyzed in this study are available in the DDBJ Sequence Read Archive under accession numbers DRR024544– DRR024549. The numerical data underlying Fig. 3 and Supplementary Fig. S3 are provided in Supplementary Tables S7 and S8, respectively.

### Author contributions

1. H. U.: Formal analysis, Investigation, Writing—original draft. K. T.: Investigation. S. N.: Formal analysis. S. I.: Funding acquisition, Methodology, Supervision. A. H.: Conceptualization, Data curation, Formal analysis, Funding acquisition, Investigation, Methodology, Supervision, Visualization, Writing—original draft, Writing—review & editing.

## ACKNOWLEDGMENTS

The authors thank Eiji Nitasaka for providing the *I. nil* seeds and Nagisa Okuda for taking the photograph shown in Fig. 1B. The authors also thank Jun Minagawa and Chiyo Noda for their guidance on pigment analysis, Kazuyo Ito, Tomoyo Takeuchi, Kiyoko Kuzunishi, and Naoko Koyama for their technical assistance, and the Model Organisms Facility and the Trans-Omics Facility of the NIBB Trans-Scale Biology Center for their support.

