## Supplementary Figures S1-S3 for "Disruption of the single-copy GOLDEN2-like gene underlies the classical *yellow* and *yellow-mutable* mutations of Japanese morning glory"

A

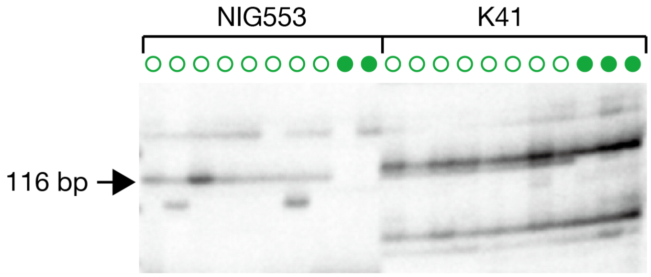


C

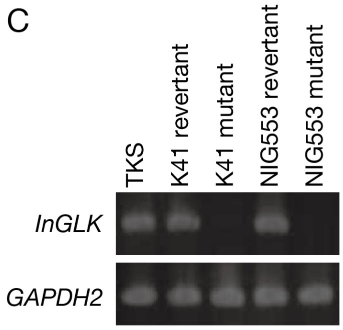


B

```
         Tpn12
         ▼
1        actttaattt gtatcagTCA AAGGAGAGTA TAGACGCGGC TATTGGAGAT        50
                              S   K   E   S   I   D   A   A   I   G   D

                                               BfaI
51       GTTTTATCGA AGCCATGGCT GCCACTTCCG CTAG                          84
         V   L   S   K   P   W   L   P   L   P   L
```

**Supplementary Fig. S1. Molecular characterization of *InGLK* in *yellow-mutable* and germinal revertant lines.**

(A) Simplified transposon display analysis of genomic DNA fragments adjacent to *Tpn1*-family transposons. The *yellow-mutable* lines NIG553 and K41 and their germinal revertants were analyzed using genomic DNA digested with *Bfa*I. Open and filled green circles indicate mutant and revertant plants, respectively. The arrowhead indicates the 116-bp band that co-segregated with the *yellow-mutable* phenotype.

(B) Sequence of the genomic fragment isolated by simplified transposon display. Putative exon and intron sequences are shown in uppercase and lowercase letters, respectively. The deduced amino acid sequence is shown below the nucleotide sequence. The target site sequence of *Tpn12* is underlined. Primer-derived sequences were omitted, resulting in an 84-bp displayed sequence from the 116-bp fragment.

(C) RT-PCR analysis of *InGLK* expression in young leaves of the K41 and NIG553 lines. Primers for *InGLK* were designed in the exon downstream of the *Tpn12* insertion site. *GAPDH2* was used as an internal control. PCR amplification was performed for 30 cycles.

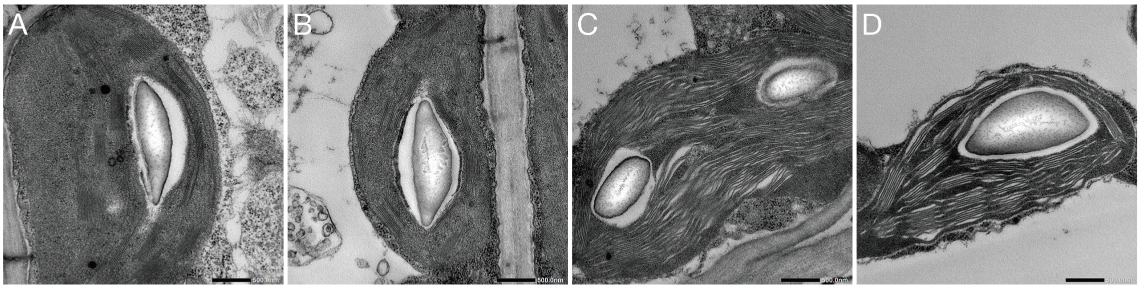


**Supplementary Fig. S2. Chloroplast ultrastructure in a green somatic reversion sector and the corresponding yellowish-green background area of a *yellow-mutable* leaf.** Transmission electron microscopy was used to examine chloroplasts in palisade mesophyll cells (A, B) and spongy mesophyll cells (C, D). Images from the green somatic reversion sector are shown in A and C, and those from the corresponding yellowish-green background area are shown in B and D. Scale bars: 500 nm.

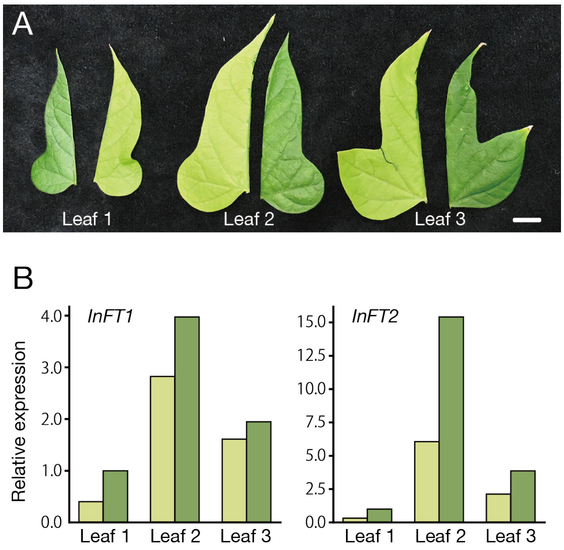


**Supplementary Fig. S3. *InFT1* and *InFT2* expression in green somatic reversion sectors and yellowish-green background areas of *yellow-mutable* leaves.**

(A) Photographs of the three true leaves used for RT-qPCR analysis. The leaves are designated Leaves 1–3 from left to right. Each leaf was divided along the midrib into a green reversion sector and the corresponding yellowish-green background area. Scale bar: 10 mm. (B) Relative transcript levels of the FT homologs *InFT1* and *InFT2* in the areas shown in (A). Expression levels were normalized to *gATPase* and are shown relative to the green reversion sector of Leaf 1 for each gene. Dark green and light green bars indicate green reversion sectors and yellowish-green background areas, respectively. Technical triplicates were averaged for each sample. Each pair of bars represents the two areas from a single leaf.
